# Screening Mycobacterium tuberculosis secreted proteins identifies Mpt64 as eukaryotic membrane-binding virulence factor

**DOI:** 10.1101/402099

**Authors:** Chelsea E. Stamm, Breanna L. Pasko, Sujittra Chaisavaneeyakorn, Luis H. Franco, Vidhya R. Nair, Bethany A. Weigele, Neal M. Alto, Michael U. Shiloh

## Abstract

*Mycobacterium tuberculosis* (Mtb), the causative agent of tuberculosis, is one of the most successful human pathogens. One reason for its success is that Mtb can reside within host macrophages, a cell type that normally functions to phagocytose and destroy infectious bacteria. However, Mtb is able to evade macrophage defenses in order to survive for prolonged periods of time. Many intracellular pathogens secret virulence factors targeting host membranes and organelles to remodel their intracellular environmental niche. We hypothesized that Mtb exported proteins that target host membranes are vital for Mtb to adapt to and manipulate the host environment for survival. Thus, we characterized 200 exported proteins from Mtb for their ability to associate with eukaryotic membranes using a unique temperature sensitive yeast screen and to manipulate host trafficking pathways using a modified inducible secretion screen. We identified five Mtb exported proteins that both associated with eukaryotic membranes and altered the host secretory pathway. One of these secreted proteins, Mpt64, localized to the endoplasmic reticulum during Mtb infection of murine and human macrophages and was necessary for Mtb survival in primary human macrophages. These data highlight the importance of exported proteins in Mtb pathogenesis and provide a basis for further investigation into their molecular mechanisms.

**Importance:** Advances have been made to identify exported proteins of *Mycobacterium tuberculosis* during animal infections. These data, combined with transposon screens identifying genes important for *M. tuberculosis* virulence, have generated a vast resource of potential *M. tuberculosis* virulence proteins. However, the function of many of these proteins in *M. tuberculosis* pathogenesis remains elusive. We have integrated three cell biological screens to characterize nearly 200 *M. tuberculosis* exported proteins for eukaryotic membrane binding, host subcellular localization and interactions with host vesicular trafficking. In addition, we observed the localization of one exported protein, Mpt64, during *M. tuberculosis* infection of macrophages. Interestingly, although Mpt64 is exported by the Sec pathway, its delivery into host cells was dependent upon the action of the Type VII Secretion System. Finally, we observed that Mpt64 contributes to the virulence of *M. tuberculosis* during infection of primary human macrophages.

## Introduction

Tuberculosis caused by *Mycobacterium tuberculosis* (Mtb) is a persistent, global epidemic. While the number of deaths due to Mtb fell below 2 million in 2015, there were over 9 million new cases (1) and the incidence of multidrug-resistant Mtb is increasing (1), highlighting the need for new anti-tuberculosis therapies. In addition, the only currently available vaccine, *Mycobacterium bovis* Bacille-Calmette-Guérin (BCG), is ineffective in preventing pulmonary tuberculosis infection (2). Thus, understanding the intracellular survival mechanisms employed by Mtb is vital to developing new anti-tuberculosis treatments and vaccines.

Macrophages, phagocytic innate immune cells that are generally competent for bacterial killing, represent the primary intracellular niche for Mtb. Some of the antimicrobial mechanisms utilized by macrophages include acidification of the phagosome, production of reactive oxygen and nitrogen species, fusion of lysosomes to bacterial containing phagosomes and autophagy (3-6). However, despite these robust defenses, Mtb survives inside macrophages during its infectious life cycle. To facilitate its survival Mtb has evolved to resist macrophage defenses, either by directly protecting the bacterial cell from damage (7-9) or by modulating the macrophage’s ability to shuttle the bacteria through the traditional phagolysosomal maturation process (10). In that way, Mtb prevents its intracellular compartment from acidifying (11) and fusing (12) with the destructive lysosome. Genetic studies have identified several Mtb proteins important for remodeling host membrane trafficking (13-15). For example, Mtb *Rv3310* encodes SapM, a secreted acid phosphatase (16) that converts phosphatidylinositol 3-phosphate (PI3P) to phosphatidylinositol. Loss of PI3P from the phagosome membrane is sufficient to prevent fusion of phagosomes with late endosomes (17, 18). Importantly, many genes reported to be important for Mtb survival inside macrophages remain uncharacterized (13, 14, 19) and the manipulation of the host cell by Mtb remains poorly understood.

The problem of intracellular survival faced by Mtb is also shared by other bacterial pathogens, and many of these organisms utilize specialized secretion systems to deliver molecules into the host cell to establish a unique intracellular niche (20). For example, some Gram-negative pathogens use needle-like machines that span the bacterial and host cell membranes to inject protein cargo into the host (21-24). Encoded by a Type III Secretion System (TTSS), the needle-like machine shares its structure with the bacterial flagellum and is expressed by a number of intracellular pathogens including *Salmonella enterica* subsp. *enterica* serovar Typhimurium, *Shigella flexneri*, and *Chlamydia trachomatis* (21, 22). Another specialized secretion machine called a Type IV Secretion System is found in Gram-positive and Gram-negative bacteria and can used for translocation of nucleic acid through conjugation (23) while in many human and plant pathogens such as *Legionella pneumophila, Coxiella burnetii* and *Agrobacterium tumefaciens,* it also transports effector proteins that promote bacterial survival (25, 26). The Type VI Secretion System is structurally analogous to the bacteriophage tail spike and is expressed by pathogenic bacteria including *Francisella tularensis, Pseudomonas aeruginosa and Vibrio cholera* (24, 27, 28). Finally, Mtb encodes multiple Type VII secretion systems, discussed below, that are important in pathogenesis (29, 30). However, the identity and functions of the Type VII dependent secreted proteins remain unknown. Despite structural differences, a common theme is that these elaborate secretion systems aid in pathogenesis by delivering virulence proteins called “effectors” to the host cell.

Effectors are proteins that promote bacterial survival by manipulating vital cellular processes including signal transduction, vesicular trafficking and the cytoskeleton (31-33). Like the secretion systems themselves, the repertoire of effectors expressed by each pathogen can differ, adapted specifically for each unique life cycle. However, a major common target for effectors are host membranes. For example, SifA from *S. typhimurium* is prenylated inside the host cell and localizes to the Salmonella containing vacuole (SCV) (34). SifA recruits lysosomes to maintain SCV membrane integrity and its membrane interaction is vital to Salmonella pathogenicity (34, 35). The *Legionella pneumophila* Type IV secreted effector SidM is anchored to the Legionella containing vacuole and disrupts host vesicle trafficking by sequestering and modifying Rab1 (36). Some effectors can also function by directly modifying membranes such as IpgD from *S. flexneri* that hydrolyzes phosphatidylinositol 4,5-bisphosphate (PI(4,5)P_2_) to phosphatidylinositol 5-monophosphate (PI5P) leading to membrane blebbing and bacterial uptake into host cells (37). Thus, both host membranes themselves and membrane-dependent processes represent valuable targets for bacterial effectors (31, 33, 38) as we recently showed for a variety of bacterial pathogens (39).

The virulence functions of most Mtb secreted proteins are poorly understood. Although Mtb does not encode a T3SS, it does contain the conserved general secretion systems Sec and Tat (40), as well as multiple Type VII (also called ESX) secretion systems (30). The ESX-1 system is required for Mtb virulence in macrophages and mouse models of infection (29) through modulation of immune functions such as induction of Type I interferon via the secreted effectors EsxA and EsxB (41, 42). Similarly, EsxH, a substrate of the ESX-3 system, interacts with a component of the host endosomal sorting complex required for transport (ESCRT) machinery, leading to decreased co-localization of Mtb with lysosomes (43) and inhibition of antigen presentation in infected macrophages (44). Substrates of the accessory SecA2 system such as the protein kinase PknG and the esterase LipO are also important for Mtb virulence by contributing to phagosome maturation arrest (PMA) (13, 18, 45, 46). Thus, because membrane processes are high value targets of many bacterial effectors (39, 47), and Mtb has a large repertoire of exported proteins of unknown function (48-52), we hypothesized that some of the Mtb exported proteins are membrane-binding effectors with virulence activities.

To test our hypothesis we generated a library of 200 exported proteins from Mtb, tested whether they individually bound yeast membranes in a life or death assay, and characterized their ability to alter host protein secretion in an inducible secretion assay. We also determined the subcellular localization of each membrane-binding protein using fluorescence microscopy. By combining data from the cell biological screens, we identified five Mtb exported proteins that localized to eukaryotic membranes and inhibited protein transport through the host secretory pathway. One protein, Mpt64 (Rv1980c), localized to the ER during both heterologous expression in HeLa cells and Mtb infection of macrophages. Though Mpt64 is a Sec substrate, its access to the macrophage cytoplasm was dependent on the ESX-1 secretion system. Finally, bacteria lacking Mpt64 were attenuated for growth in primary human macrophages, demonstrating an important virulence activity for Mpt64.

## Results

### Categorization of putative effector-like proteins from Mtb

Through the analysis of published datasets we identified Mtb proteins that may function as secreted effectors (Supplemental Table 1). For clarity, we define these proteins as <u>M</u>ycobacterial <u>e</u>xported <u>p</u>roteins (MEPs), as this encompasses proteins that are secreted as well as those delivered to the mycobacterial surface, which can still modulate interactions with the host (53, 54). We used the following criteria to assemble a library of MEPs: 1) Mtb secreted proteins identified via proteomic approaches (49-52, 55, 56), 2) Mtb proteins known to be involved in manipulation of host vesicular trafficking pathways, such as ones that induce mammalian cell entry (MCE) (57-59) or phagosome maturation arrest (13, 14, 60), 3) a subset of PE/PPE proteins and proteins related to those encoded by Type VII (ESX-1) loci (61-63), and 4) proteins involved in virulence, ranging from defined to unknown functions (19, 54, 64-66). We then used Gateway recombination cloning to subclone MEPs from the freely available Mtb ORFome Gateway compatible library (BEI) into destination vectors for a variety of subsequent assays. The comprehensive list of MEPs is shown in Supplemental Table 1.

### Mtb encodes exported proteins that interact with eukaryotic membranes

To identify membrane binding Mtb proteins, we used a system that leverages the signal transduction of the essential yeast GTPase Ras to promote growth and division (67). Ras is lipidated at a unique sequence called the CaaX box that promotes its localization to the plasma membrane, where it can be activated by Cdc25, a guanine nucleotide exchange factor (67, 68). In a yeast strain with a temperature sensitive *CDC25* allele, yeast can only grow at the permissive temperature (25°C) but not the restrictive temperature (37°C) because Ras activation requires interaction with Cdc25. Heterologous expression of a non-lipidated, constitutively-active Ras whose activity is independent of Cdc25 (Ras^mut^) can rescue yeast growth at the restrictive temperature when Ras is recruited to intracellular membranes by fusion to a membrane binding protein (Fig. 1A). This system has been used to successfully identify membrane binding effectors from Gram negative pathogens (39). To identify membrane-localizing proteins from Mtb, we subcloned MEPs into a destination vector for yeast expression that generates an in-frame fusion of the MEP to Ras^mut^. We transformed *S. cerevisiae cdc25^ts^* (39, 67) individually with each of the 200 MEP fused to Ras^mut^ and incubated them at both permissive and restrictive temperatures (Figure 1B). We identified 52 Mtb proteins that rescued *S. cerevisiae cdc25^ts^* growth at the restrictive temperature (Fig. 1B, C).

**Figure 1.**
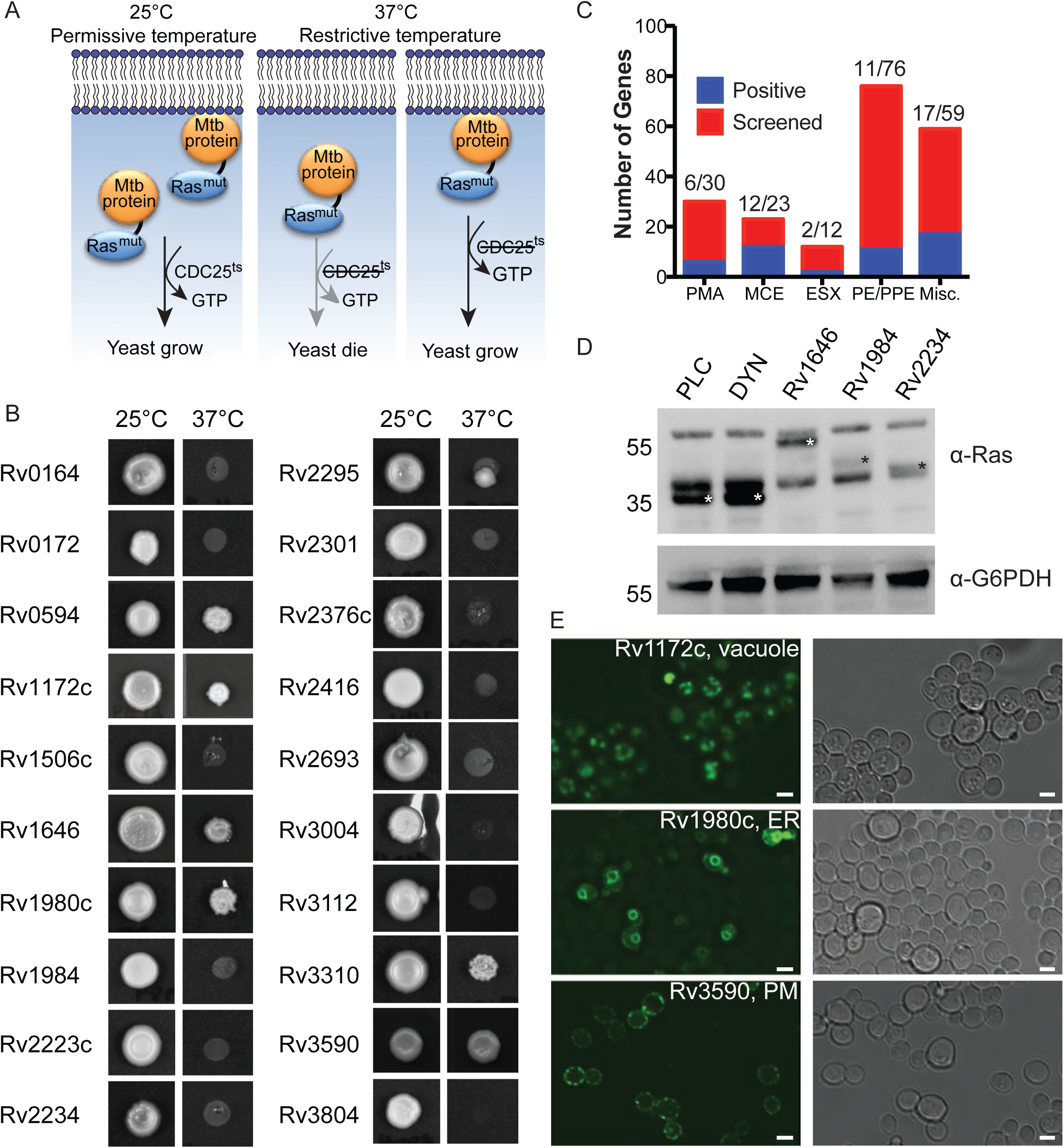
*M. tuberculosis* exported proteins interact with yeast membranes. (**A**) Ras rescue assay schematic. (**B**) *S. cerevisiae* (cdc25^ts^) transformed with Mtb protein fusions to Ras^mut^, duplicate plated and incubated 48-72 h at the permissive (25°C) and restrictive (37°C) temperatures. Shown are representative images of 20 yeast strains. (**C**) Summary results of Ras rescue screen. (**D**) Western blot of lysates from yeast transformed with the indicated fusion proteins and probed with anti-Ras or anti-G6PDH antibodies. Ras^mut^ fusion proteins are marked by white or black asterisks. PLC (phospholipase C) and DYN (dynamin) are fusions to Ras^mut^ known to be membrane associated (PLC) or cytoplasmic (DYN). (**E**) Representative fluorescence microscopy of *S. cerevisiae* (INVSc1) transformed with GFP-MEP fusion proteins. Scale bars are 3μm.

We confirmed expression of the Ras^mut^-MEP fusion proteins by Western blotting (Figure 1D). In addition, we determined the membrane localization of each MEP by fluorescence microscopy of GFP-MEP fusion proteins in yeast (Fig. 1E). It has been established that Ras can function from membranes other than the plasma membrane (69, 70) and Ras^mut^ maintains this function (39). Thus, using fluorescence microscopy we observed GFP-MEP fusion proteins localizing to distinct subcellular compartments including vacuoles, ER and plasma membrane (Figure 1E). Together these results show that 25% of the MEPs tested could associate with the membranes of a variety of organelles in *S. cerevisiae*.

### Subcellular localization of membrane-localizing MEPs

While many cellular processes are conserved in eukaryotes, humans represent the primary natural host for Mtb. Therefore, to confirm that MEPs that rescued *S. cerevisiae cdc25^ts^* growth at 37°C also bound mammalian membranes and to determine their subcellular localization in human cells, we transiently transfected HeLa cells with vectors for constitutive expression of GFP-MEP fusion proteins and then used fluorescence microscopy with co-localization markers to identify the specific membrane to which each MEP localized (Figure 2A). We identified GFP-MEP fusion proteins that localized to a variety of subcellular compartments including the ER, Golgi, mitochondria and peroxisomes (Figure 2A and 2B). The largest proportion of the GFP-MEP fusion proteins expressed in human cells co-localized with the ER marker calreticulin (Figure 2B). In yeast, we observed a similar number of GFP-Mtb fusion proteins which localized to compact, punctate structures (data not shown). Although there was only moderate overlap in the subcellular localization identified between yeast and HeLa cells (Figure 2B), we were able to verify that the proteins identified by the Ras rescue assay are localized to membranes in human cells.

**Figure 2.**
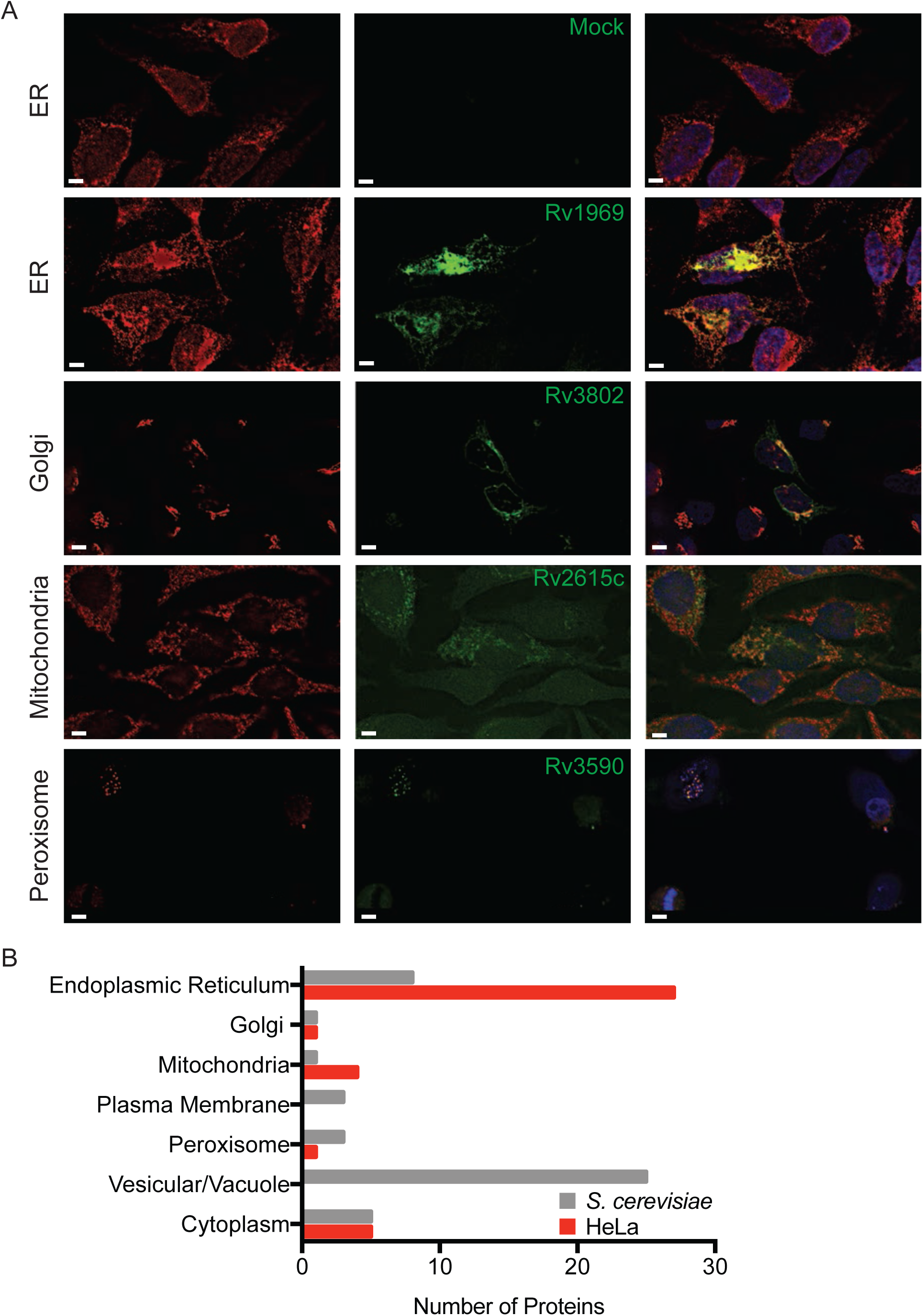
Host subcellular localization of membrane-binding MEP. (**A**) Representative fluorescent images of HeLa cells transfected with the indicated GFP-MEP fusion proteins (green) and stained with antibodies (red) to calreticulin (ER), GM-130 (Golgi), Tom20 (Mitochondria) or PMP70 (Peroxisomes). Scale bars are 5μm. (**B**) Comparison of the organelle localization of MEP expressed in yeast and HeLa cells.

### A subset of mycobacterial exported proteins alter the host secretory pathway

Membrane-bound cargo is transported within the host cell and to the extracellular space in dynamic vesicular trafficking pathways. The membrane-bound and soluble proteins important for these processes are frequent targets of bacterial effectors (47, 71). Indeed, Mtb is known to target and manipulate trafficking pathways through incompletely understood mechanisms (43, 72, 73). To determine if Mtb proteins can broadly affect the host general secretion pathway as an indicator of interaction with membranes and intracellular vesicular trafficking, we took advantage of the reverse dimerization system (74). In this system, a secreted protein of interest is sequestered in the ER by fusion to a conditional aggregation domain (CAD). Addition of a solubilization molecule that disrupts the CAD then frees the fusion protein for trafficking and secretion into the extracellular space. We used a fusion of human growth hormone (hGH) to the CAD domain of the ligand-reversible crosslinking protein, FKBP F36M. Thus, hGH can be quantified in cell supernatants by ELISA after addition of the small molecular, D/D Solubilizer (Figure 3A) (74, 75). We transfected HeLa cells expressing hGH-CAD with each MEP individually, a negative control protein (GFP), or an enterohemorrhagic *Escherichia coli* effector (EspG) that inhibits secretion by promoting the tethering of vesicles to the Golgi apparatus (75, 76). When we treated transfected cells with D/D Solubilizer, we observed a spectrum of effects on the secretion of hGH including increased, decreased and normal hGH secretion (Figure 3B and Supplemental Table 1). Using a cut-off of normalized hGH secretion below 0.25 or above 1.75, we identified 10 proteins that decreased hGH secretion and 10 proteins that increased hGH secretion as compared to the GFP control (Figure 3C). We next compared the MEPs that altered host secretion to those that bound eukaryotic membranes, and identified five proteins with overlapping activities: Rv0594, Rv1646, Rv1810, Rv1980c and Rv2075 (Figure 3D). During expression in HeLa cells, all but one protein localized to the ER and all five proteins reduced hGH secretion (Figure 3E).

### Mpt64 N-terminus is important for membrane binding and secretion inhibition

We focused on the protein Rv1980c, also known as Mpt64, as it is a secreted protein that is highly antigenic during human tuberculosis infection (77, 78). Furthermore, Mpt64 is a component of the region of difference 2 (RD2) locus, one of the genomic regions deleted during attenuation of the *M. bovis* BCG vaccine strain (79). Loss of RD2 from Mtb attenuates its virulence, and complementation with a three gene cluster that includes Mpt64 can partially restore virulence (80).

**Figure 3.**
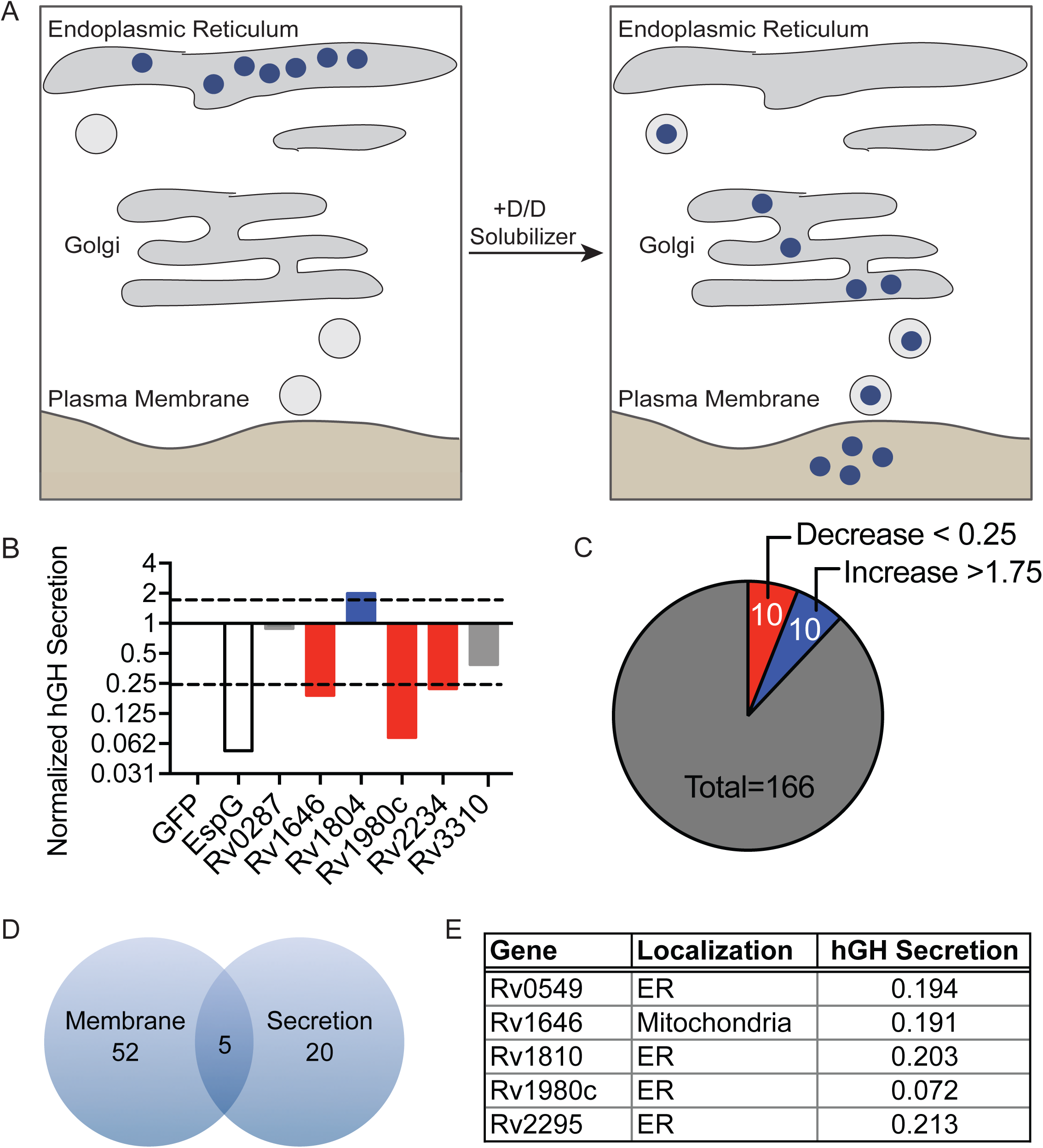
*M. tuberculosis* exported proteins alter hGH secretion (**A**) Inducible secretion assay schematic. (**B**) Supernatant hGH ELISA from HeLa cells transfected overnight with hGH-CAD and either GFP or GFP-MEP fusion proteins prior to addition of drug to allow for hGH secretion. hGH secretion by GFP-MEP transfected cells was normalized to hGH secretion of cells transfected with GFP alone. (**C**) Summary of results from inducible secretion screen. (**D**) Venn diagram of MEP that are membrane-localized, alter host secretion, or both. (**E**) Table summarizing the membrane localization and degree of hGH secretion in cells transfected with the five overlapping proteins from **D**.

Mpt64 is a 25kDa protein with a predicted signal peptidase 1 cleavage site between amino acids 23 and 24, such that the mature, secreted form of the protein starts at amino acid 24 (50, 51, 81). While the solution structure of Mpt64 was previously solved (82), the structure does not align to a known catalytic domain but does contain a domain of unknown function (DUF3298). This domain is also present in the lysozyme-binding anti sigma factor RsiV (83). Despite structural homology between Mpt64 and RsiV (Supplemental Figure 1A) (84), there is little primary sequence homology. To determine if Mpt64 binds lysoszyme, we purified recombinant Mpt64 from *E. coli* and tested binding to human or hen egg white lysozyme in an *in vitro* pull down assay (83). Using this assay we were unable to demonstrate lysozyme binding by Mpt64 (Supplemental Figure 1B). We next used the solution structure to guide truncation analysis of Mpt64 in order to identify the membrane binding sequences of Mpt64 (Figure 4A and Figure 4B). *S. cerevisiae cdc25^ts^* expressing a fusion of Ras^mut^ with either full length Mpt64, mature Mpt64 lacking its predicted signal peptide or the N-terminal half of Mpt64 also lacking the signal peptide (Mpt64_24-143) were able to grow at 37°C, whereas *S. cerevisiae cdc25^ts^* expressing Ras^mut^ fused to the C-terminal half of Mpt64 (Mpt64_144-228) could not (Figure 4C). We detected expression of Ras^mut^ fusions of full length Mpt64 and mature Mpt64 by Western blot. In contrast, we could not detect Ras^mut^ fusions of Mpt64_24-143 or Mpt64_144-228 despite the fact that the Mpt64_24-143 fusion rescued yeast growth, suggesting that expression of Mpt64_24-143 below the limit of detection by Western blot was still sufficient to rescue yeast growth (Figure 4D). However, whether the C-terminal domain plays a role in membrane binding could not be determined definitively because we were unable to demonstrate stable fusion protein expression. To test if Mpt64 could interact with lipids directly, we expressed and purified recombinant Mpt64 and Mpt64 variants from *E. coli* and tested their ability to bind unique lipid species i*n vitro* using membranes spotted with lipids. Recombinant Mpt64_24-228 bound phosphatidylinositol 4-phosphate (PI4P), phosphatidylinositol 5-phosphate (PI5P), phosphatidylinositol 4,5-bisphosphate [PI(4,5)P_2_] and phosphatidylinositol (3,4,5)-trisphosphate [PI(3,4,5)P_3_] on PIP strips membranes (Figure 4E). Similarly, recombinant Mpt64_24-143, the N-terminal portion of the protein, also bound PI4P and PI5P with additional binding to PI3P, PI(3,4)P_2_ and phosphatidylserine (Figure 4E). However, interaction with PI(4,5)P_2_ and PI(3,4,5)P_3_ was weak, suggesting that the C-terminal region of Mpt64 modifies its interactions with host phospholipids.

**Figure 4.**
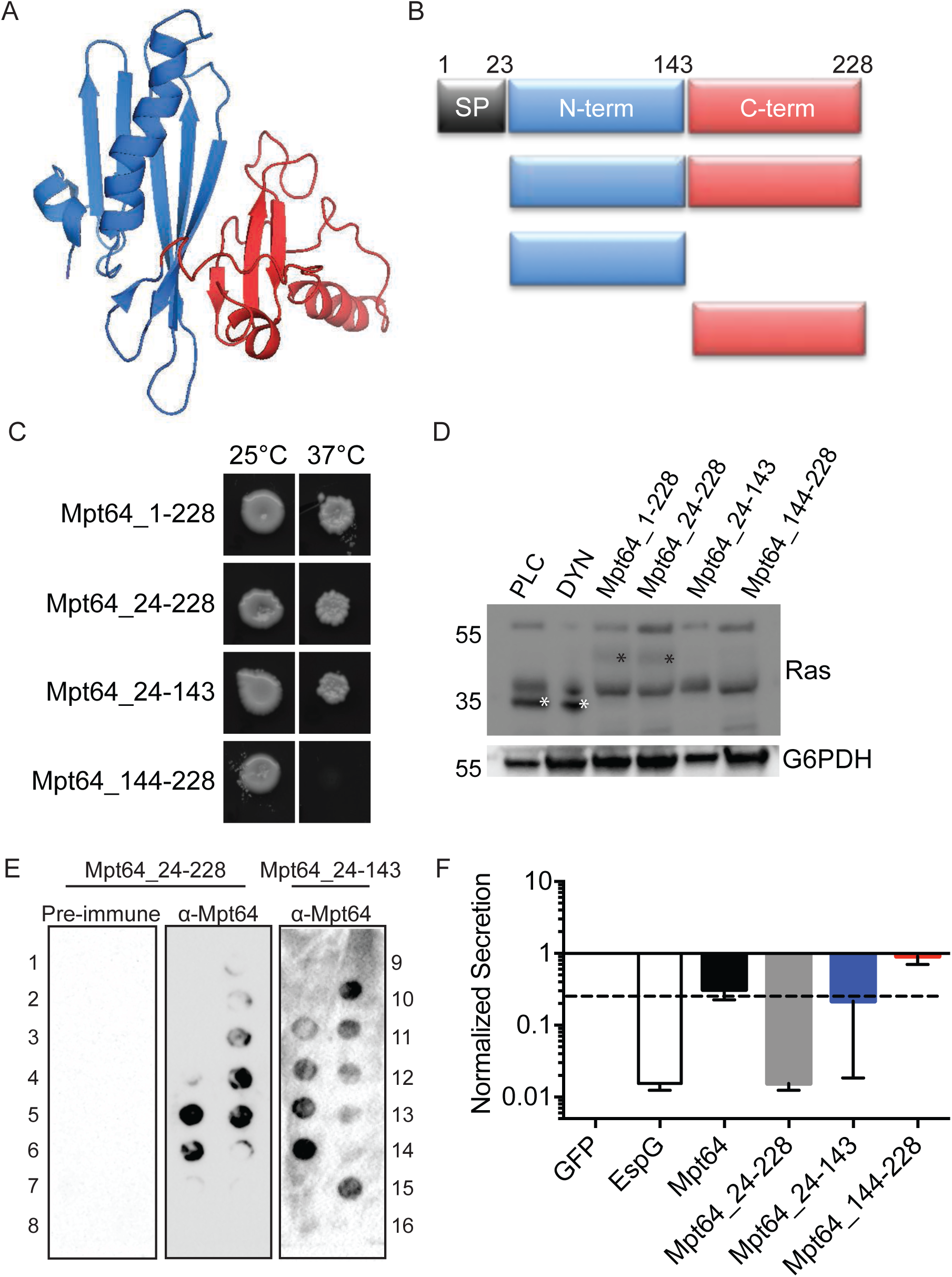
hGH secretion inhibition is dependent on membrane localization of Mpt64. (**A**) Solution structure of Mpt64. PDB 2hhi. (**B**) Schematic of Mpt64 truncations, colored to match solution structure in A. SP, signal peptide. (**C**) Full length Mpt64 or protein truncations expressed in the Ras rescue assay. (**D**) Western blot of lysates from cdc25^ts^ yeast expressing Ras^mut^ fusion proteins to Mpt64 or Mpt64 truncations. Blots were probed for anti-Ras or anti-G6PDH antibodies. Control protein bands are marked by a white asterisk and Mpt64 truncations are marked by a black asterisk. (**E**) PIP membrane strips incubated with recombinant Mpt64_24-228 or Mpt64_24-143. Binding of Mpt64 to lipids was detected by incubation with α-Mpt64 or pre-immune serum. Numbers indicate specific lipids as follows: 1, lysophosphatidic acid; 2, lysophosphatidylcholine; 3, phosphatidylinositol; 4, phosphatidylinositol-3-phopshate (PI3P); 5, PI4P; 6, PI5P; 7, phosphatidylethanolamine; 8, phosphatidylcholine; 9, sphingosine 1-phosophate; 10, phosphatidylinositol-3,4-bisphosphate [PI(3,4)P2]; 11, PI(3,5)P2; 12, PI(4,5)P2; 13, phosphatidylinositol-3,4,5-trisphosphate [PI(3,4,5)P3]; 14, phosphatidic acid; 15, phosphatidylserine; 16, blank. (**F**) ELISA results of hGH in supernatants of cells co-expressing full length Mpt64, Mpt64 truncations or controls.

Finally, we sought to determine if the N-terminal portion of Mpt64 was also sufficient to inhibit hGH secretion using the hGH-CAD assay. We co-transfected hGH-CAD expressing HeLa cells with the same Mpt64 truncations and determined their ability to inhibit hGH secretion in the presence of drug. Similar to the Ras rescue assay, full length, mature and Mpt64_24-143 inhibited hGH secretion compared to the GFP control. In contrast, co-transfection of Mpt64_144-228 with hGH-CAD had no effect on its secretion (Figure 4F). These data suggest that the ability of Mpt64 to bind membranes and to inhibit host secretion *in vitro* is dependent on the N-terminus of the protein.

### Mpt64 ER localization depends on its N-terminus

As full-length Mpt64 localized to the ER in yeast and HeLa cells (Fig. 1,3) we next tested the impact of Mpt64 truncations on ER localization. We first determined the phenotypic localization of Mpt64 truncations expressed as GFP fusions in yeast using fluorescence microscopy. Mpt64_1-228 and Mpt64_24-228 localized in a ring indicative of the ER (85, 86)(Figure 5A). In contrast Mpt64_144-228, which did not rescue yeast growth in the Ras Rescue assay (Figure 4C), was diffuse throughout the yeast cell (Figure 5A). Interestingly, Mpt64_24-143 localized to bright puncta within the cells (Figure 5A). To confirm the N-terminal dependence of Mpt64 localization, we transfected HeLa cells with GFP fusions to each Mpt64 truncation and assayed for co-localization with calreticulin through immunofluorescence microscopy. While full length Mpt64, mature Mp64 and Mpt64_24-143 co-localized with calreticulin, Mpt64_144-228 did not (Figure 5B). These results demonstrate that Mpt64 localizes to the ER during exogenous expression in both yeast and mammalian cells and the N-terminal 143 amino acids are sufficient for Mpt64 to localize to membranes.

**Figure 5.**
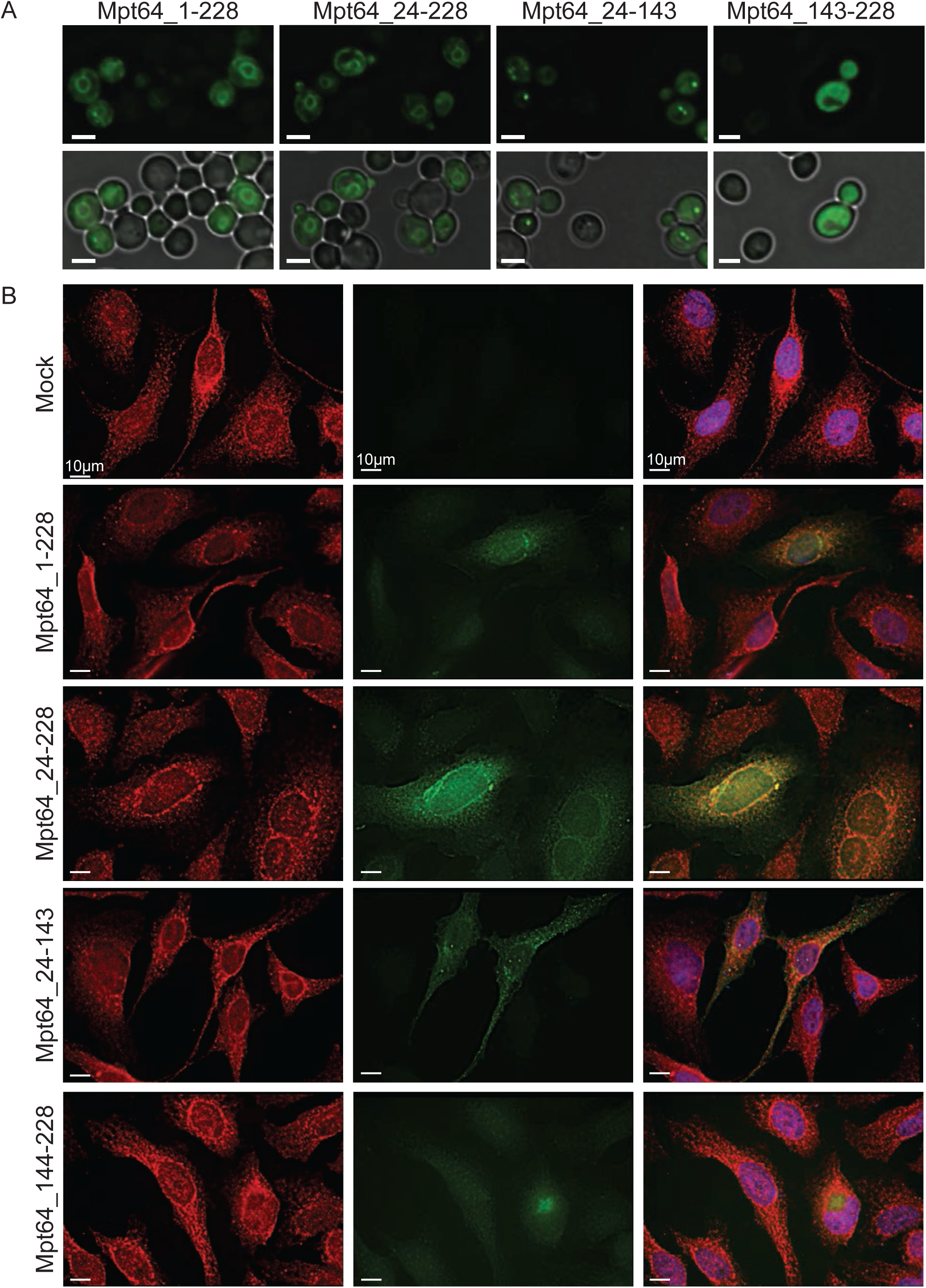
Mpt64 localizes to the endoplasmic reticulum during heterologous expression in yeast and HeLa cells. (**A**) Immunofluorescence (top panels) and bright field overlay (bottom panels) images of *S. cerevisiae* transformed with GFP fusion proteins to Mpt64 truncations. Scale bars are 3 μm. (**B**) HeLa cells transfected overnight with GFP fusion proteins (green) and stained for ER localization with anti-calreticulin antibody (red). Nuclei are stained with DAPI (blue). Scale bars are 10 μm.

### Secreted Mpt64 localizes to the ER during infection

Although we observed Mpt64 localization to the ER in yeast (Figure 1E and Figure 5A) and HeLa cells (Figure 5B), we wanted to determine if endogenous, untagged Mpt64 localizes to the ER during an Mtb infection of macrophages. To that end, we infected mouse RAW267.4 macrophages with mCherry-labeled Mtb at an MOI of 20:1 and fixed cells at various time points after infection. We then used a rabbit polyclonal antibody developed against recombinant, mature Mpt64 protein to track Mpt64 secretion from Mtb into macrophages using immunofluorescence microscopy. Of note, this antibody was generated without complete Freund’s adjuvant in order to avoid any cross-reactivity against Mtb antigens generated by the use of this adjuvant. As little as four hours after infection, endogenous Mpt64 was detected in both the cytoplasm and ER of host cells (Figure 6A, upper panels). When we performed the same experiment with MtbΔeccD1, a strain that lacks the Type VII secretion system secretion pore that cannot secret ESAT-6 (29, 87, 88) and does not result in communication between the phagosome and cytoplasm (41, 42, 89), Mpt64 appeared to be secreted but trapped adjacent to the bacteria (Figure 6A, lower panels), suggesting that it could not escape the phagosome. Importantly, Mpt64 was detected in the culture filtrate prepared from MtbΔeccD1 (Supplemental Figure 2A). Thus, although Mpt64 is likely secreted from Mtb by the canonical Sec-dependent pathway, its access to the macrophage cytoplasm and other targets in the cell was dependent on the Type VII secretion system.

**Figure 6.**
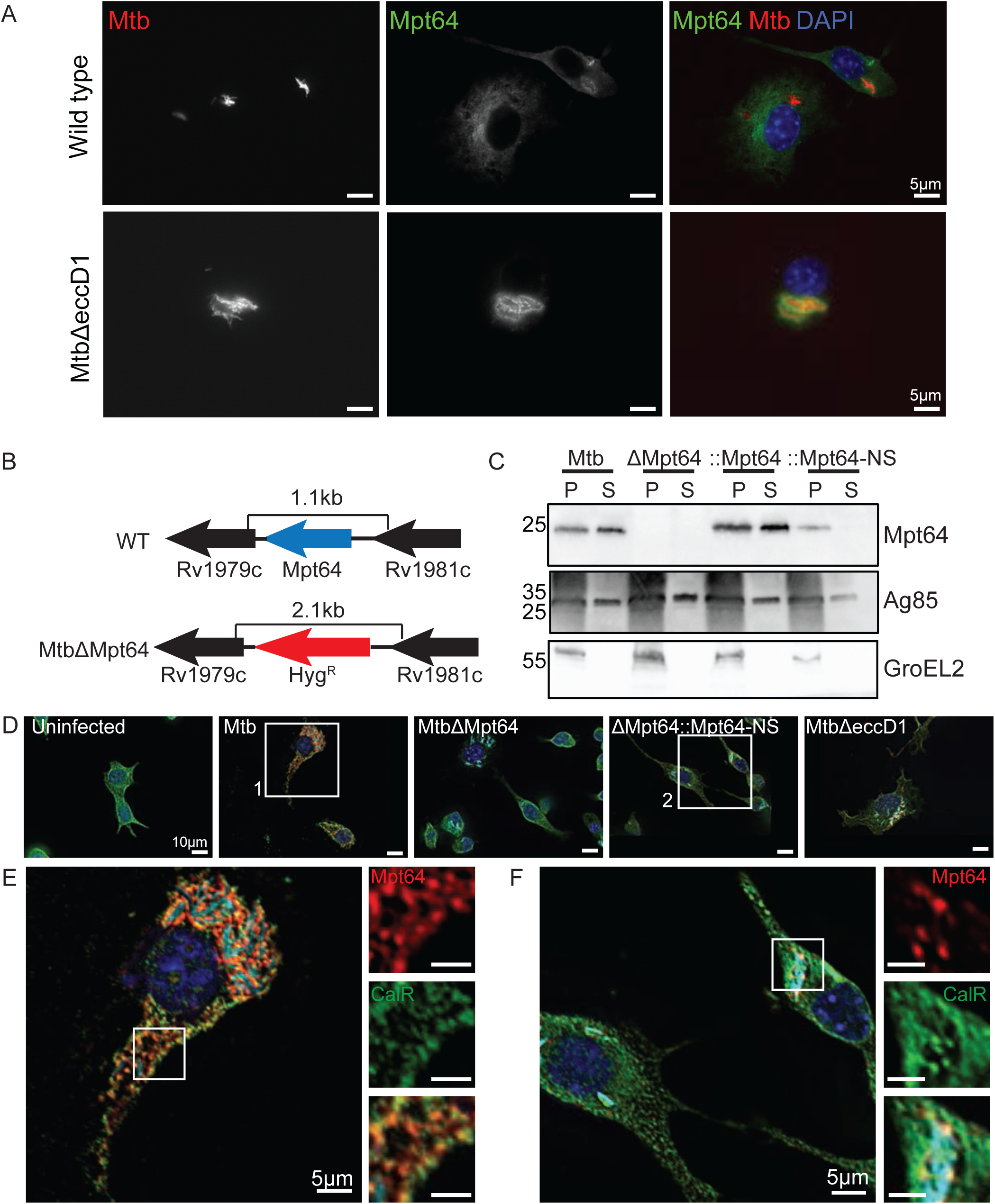
Mpt64 ER localization is ESX1-dependant during *M. tuberculosis* infection of macrophages. (**A**) RAW267.4 murine macrophages were infected with mCherry-expressing (red) WT (upper panels) or MtbΔeccD1 (lower panels) for 4 hours at an MOI 20:1. Cells were fixed and stained for Mpt64 (green) and nuclei (blue). Scale bars are 5μm. (**B**) Schematic detailing in-frame deletion of *mpt64* by insertion of a hygromycin resistance gene. (**C**) Western blot detecting expression of Mpt64, Ag85 and GroEL2 in either the lysate of the cell pellet (P) or culture supernatant (S) of four Mtb strains. (**D**) RAW267.4 macrophages were infected with the indicated strains of mCherry-expressing (cyan) Mtb for 4 hours at an MOI 20:1. Cells were fixed and stained for Mpt64 (red), calreticulin (green) and nuclei (blue). Images in (**E**) and (**F**) correspond to box 1 and box 2, respectively. Scale bars are 10μm. (**E**) Enlarged image from box 1 in (**D**) of an Mtb-infected macrophage stained for Mpt64, calreticulin and DAPI. Insets show an area of Mpt64-calreticulin co-localization. Scale bars are 5μm. (**F**) Enlarged image from box 2 in (**D**) of macrophages infected with MtbΔMpt64::Mpt64-NS and stained for Mpt64, calreticulin and DAPI. Scale bars are 5 μm.

In order to better understand the role of Mpt64 in Mtb virulence, we used mycobacteriophage (90-92) to introduce the hygromycin resistance cassette into the *Mpt64* gene to create an in-frame deletion (Figure 6B). We confirmed disruption of *Mpt64* by PCR (Supplemental Figure 2B) and loss of Mpt64 by the absence of protein on Western blot (Figure 6C). We then complemented MtbΔMpt64 with either full length Mpt64 (MtbΔMpt64::Mpt64) or Mpt64 lacking its signal peptide (MtbΔMpt64::Mpt64-NS) under the control of the constitutive mycobacterial strong promoter (93). Both complemented strains expressed Mpt64 but only full-length Mpt64 (MtbΔMpt64::Mpt64) could be detected in the supernatant of cultures, confirming that deletion of the signal peptide inhibits Mpt64 secretion from Mtb (Figure 6C). Furthermore, the MtbΔMpt64::Mpt64 strain had modestly higher expression of Mpt64 compared to wild-type Mtb by western blot, consistent with our use of a strong constitutive promoter for complementation. All four strains grew equally under axenic growth conditions (Supplemental Figure 2C), and we confirmed that both Mtb and MtbΔMpt64 produced phthiocerol dimycocerosate by mass spectrometry (Supplemental Figure 2D).

To test if secreted Mpt64 interacts with the ER during infection, we assessed its colocalization with calreticulin in RAW267.4 cells using confocal immunofluorescence microscopy. When we infected RAW267.4 macrophages, the Mpt64 signal in Mtb infected macrophages co-localized with calreticulin, confirming the subcellular localization of Mpt64 secreted during infection (Figure 6D,E and Supplemental Figure 3). However this co-localization was lost in cells infected with MtbΔMpt64::Mpt64-NS bacteria (Figure 6D,F and Supplemental Figure 3). As a control for antibody specificity, no Mpt64 was detected in macrophages infected with MtbΔMpt64 mutant bacteria (Figure 6D and Supplemental Figure 3). From these data, we can confirm that the signal peptide of Mpt64 is sufficient for the protein’s secretion in vivo and is required (with concerted action of the Type VII secretion system) for Mpt64 to interact with the ER during infection.

### Mpt64 contributes to early Mtb growth after aerosol infection of mice

Because Mpt64 is part of the Mtb RD2 locus that partially accounts for the attenuation of Mtb (80), and our data indicating that Mpt64 may function as a secreted effector, we investigated the role of Mpt64 in Mtb virulence in a murine model of infection. We infected BALB/c mice via aerosol with a low-dose of bacteria (∼100 CFU Mtb) and collected lungs at various time points to determine CFU and histopathology. We compared the infections of four strains: wild type Mtb, MtbΔMpt64, MtbΔMpt64::Mpt64 and MtbΔMpt64::Mpt64-NS (described in Figure 6). While all mice received equal numbers of bacteria between the four strains at day 0 (data not shown), there were fewer Mtb isolated from lungs of mice infected with the MtbΔMpt64 mutant compared to wild type (mean CFU wild type Mtb 2.7×10^6^ vs MtbΔMpt64 1.7×10^6^, p=0.07) at 21 days post infection (dpi) but no statistically significant difference by 42dpi (mean CFU wild type Mtb 5.0×10^5^ vs MtbΔMpt64 3.4×10^5^). However, by 42dpi there was a statistically significant decrease in the CFU isolated from lungs of mice infected with MtbΔMpt64::Mpt64-NS (Figure 7A). At these time points, we observed a reduction in the area of inflammation in hematoxylin and eosin (H&E) stained lungs of mice infected with MtbΔMpt64::Mpt64-NS compared to wild type. (Figure 7B and 7D). Despite modest reductions in CFU in mutant bacteria, there were no differences in mouse survival comparing the four strains (Figure 7C). Thus, although deletion of *Mpt64* entirely (MtbΔMpt64) and disruption of Mpt64 secretion (MtbΔMpt64::Mpt64-NS) resulted in a modest decrease in bacterial growth in the lungs of mice, loss of Mpt64 was not sufficient to explain the attenuation of the RD2 mutant (80).

**Figure 7.**
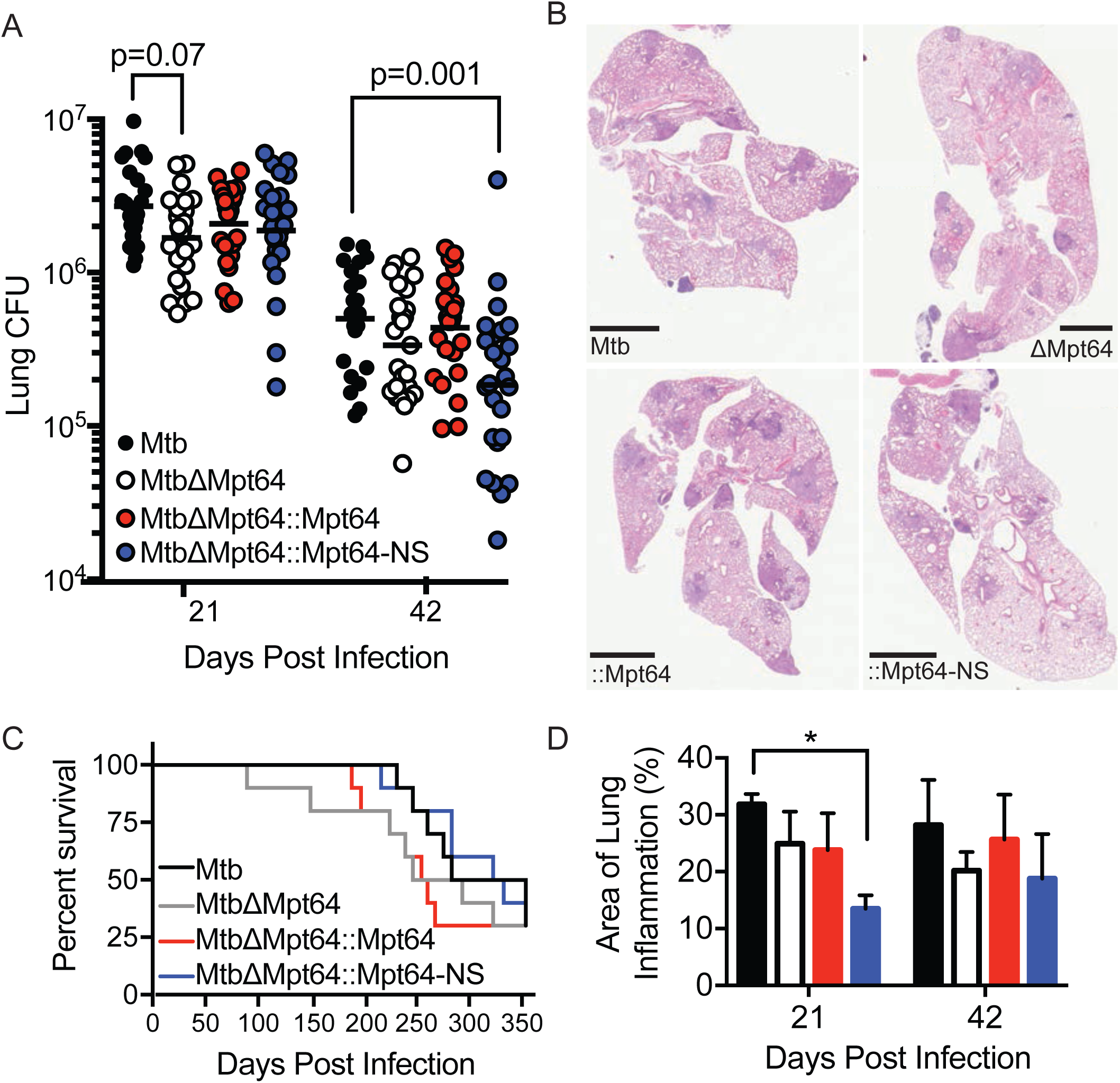
Mpt64 contributes to early Mtb growth after aerosol infection of mice (**A**) Bacterial burden in lungs of mice 21 and 42 days after aerosol infection with indicated strains of Mtb. Results are a combination of three independent experiments, n=25 mice total per group. Horizontal bar indicates the geometric mean. p values are determined by nonparametric Kruskal-Wallis analysis. (**B**) Representative images of H&E stained lungs at 42 days post-infection with the indicated strains of Mtb. Scale bars, 2mm. (**C**) Ten mice per group were monitored for survival. There were no significant differences in survival rates between groups by Kaplan–Meier analysis. (**D**) Quantitation of lung inflammation of mice infected with indicated Mtb strains. Measurement was determined using ImageJ software (NIH). Bars are colored as in (**A**). Results are the mean ±SEM for three animals per group. *p<0.02 by Kruskal-Wallis test.

### Mtb survival in human macrophages requires Mpt64

Next, we assessed whether the localization of Mpt64 in human cells is similar to that in murine macrophages. To that end, we infected primary human monocyte-derived macrophages with mCherry-expressing WT Mtb or MtbΔeccD1 and stained for Mpt64. Consistent with our data in RAW267.4 cells (Figure 6A) the secretion of Mpt64 to extra-phagosomal sites in primary human macrophages was dependent on the Type VII Secretion System (Figure 8A). Then we infected primary human macrophages with WT Mtb, MtbΔMpt64, MtbΔMpt64::Mpt64 or MtbΔMpt64::Mpt64-NS and determined the co-localization of Mpt64 with calreticulin by fluorescence microscopy (Figure 8B and Supplemental Figure 4). At 4hpi, we detected co-localization of Mpt64 with calreticulin in cells infected with WT Mtb and MtbΔMpt64::Mpt64 (top panels) but not in cells infected with MtbΔMpt64 or MtbΔMpt64::Mpt64-NS (bottom panels, Figure 8B).

**Figure 8.**
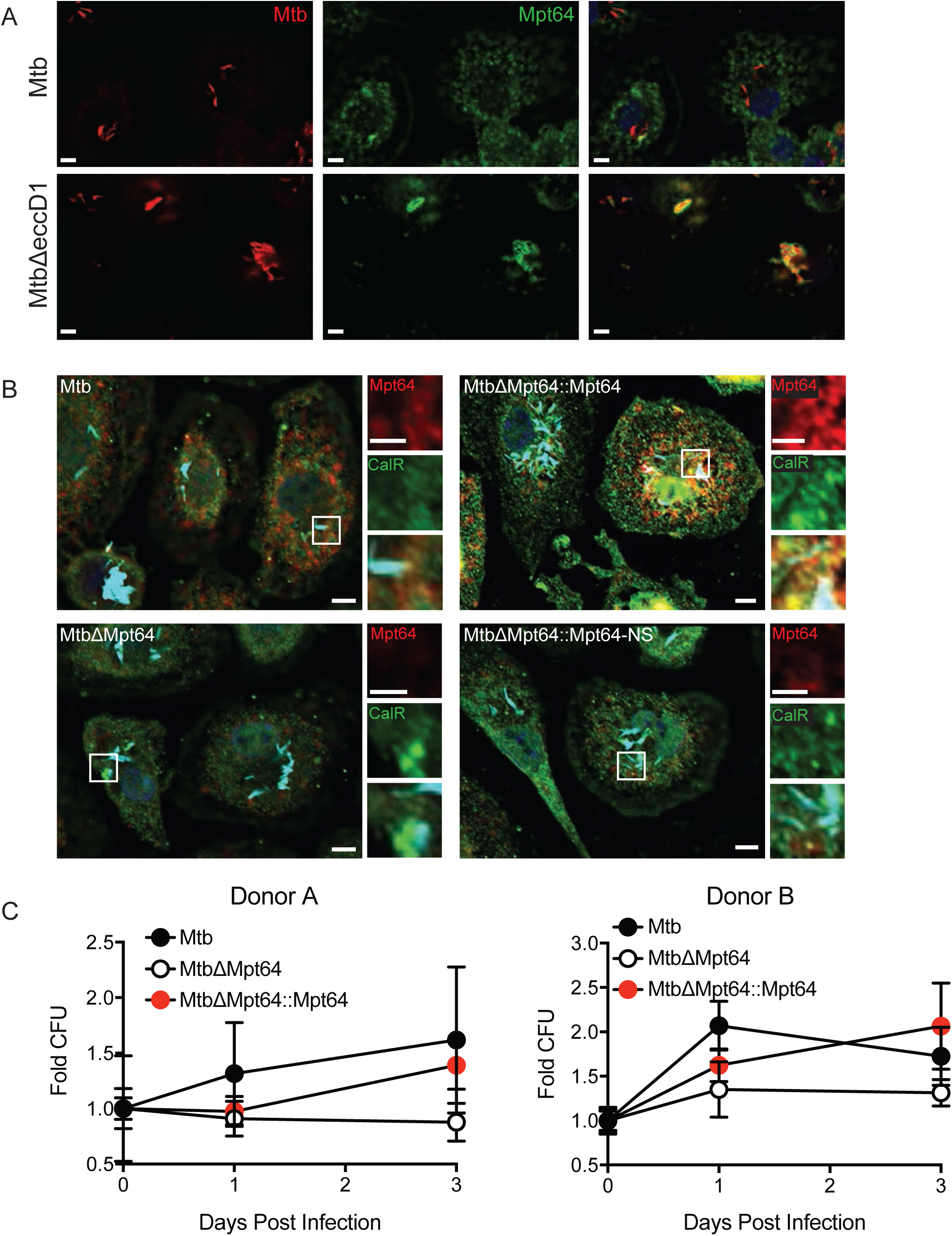
Localization and activity of Mpt64 in human macrophages (**A**) Primary human monocyte-derived macrophages were infected with mCherry-expressing (red) Mtb or MtbΔeccD1 for 4 hours at an MOI 10:1. Cells were fixed and stained for Mpt64 (green). Scale bars are 5μm. (**B**) Primary human monocyte-derived macrophages were infected with the indicated strains of mCherry-expressing Mtb for 4 hours at an MOI 10:1. Cells were fixed and stained for Mpt64 (red) and calreticulin (green). Nuclei are stained blue. Scale bars are 5 μm and scale bars of inset images are 2.5 μm. (**C**) CFU recovered from primary human macrophages from two separate donors infected with Mtb, MtbΔMpt64, or MtbΔMpt64::Mpt64 at indicated time points. CFU was normalized to the bacterial burden at day 0 and is represented as the mean with standard error. Data are representative of 6 independent experiments.

To better understand the contribution of Mpt64 in the context of human Mtb infection, we determined the growth of wild type, MtbΔMpt64 or MtbΔMpt64::Mpt64 in primary monocyte-derived human macrophages. We recovered CFU from cells directly after infection (Day 0) and one and three days post infection. While there was no difference in the CFU isolated from cells infected with each strain after one day of infection, the MtbΔMpt64 mutant was attenuated for growth three days post infection compared to wild type (Figure 8C). This growth defect was rescued in the complemented strain. Together these data indicate a role for Mpt64 in the virulence of Mtb during acute infection of human macrophages.

## Discussion

Numerous efforts have been undertaken to identify Mtb exported proteins, from lipoproteins that are incorporated into the cell wall, to virulence factors that reach the extracellular environment such as ESAT-6 (48-50). However, little is known about the function of this “secretome” as a whole. Here we took a systematic approach towards characterizing host-dependent interactions of a collated list of putative effector molecules. We created a library of 200 putative mycobacterial exported proteins and then through a series of cell biological screens characterized these MEPs for their ability to bind eukaryotic membranes, their subcellular localization and their ability to modulate secretion of a model substrate. In addition, we demonstrate that one secreted protein, Mpt64, localized to the ER during infection and was important for virulence of Mtb in human macrophages. The cohort of 200 MEPs we generated was large but not necessarily exhaustive (Supplemental Table 1). For example, the 76 PE/PPE genes we included represent less than half (45%) of the total number of PE/PPE genes in the Mtb genome (94). In addition, a recent technology called EXIT identified 593 Mtb proteins secreted during intravenous infection of mice including 38 proteins that are significantly enriched only during *in vivo* infection as compared to growth on 7H10 agar, suggesting a virulence function for these proteins (95). Of the 200 MEPs we characterized, 51 overlap with those identified by EXIT and of the 51 overlapping proteins, 25 are membrane associated in our study. This emphasizes that host membranes can be targets of Mtb virulence proteins.

We found 52 Mtb proteins that associated with eukaryotic membranes, representing nearly 25% of the total screened. When the membrane association of type III and type IV effectors from several Gram negative pathogens was explored, about 30% of effectors screened also associated with eukaryotic membranes (39). While our data are in agreement with this value, pathogens that replicate intracellularly in vacuoles had even higher numbers of membrane-associated effectors (39). This suggests that there may be additional secreted virulence proteins from Mtb that associate with the host membranes than our screen identified. Indeed, while we corroborated previously known membrane-interacting proteins such as the SecA2-secreted PI3P phosphatase SapM (17, 18), the Rac1-binding protein Ndk (96) and the cholesterol-binding Mce4A (97), we failed to identify others such as LipY which hydrolyzes extracellular lipids (98) and the ESX1 substrate ESAT-6 which interacts with the phagosomal membrane (99, 100).

GFP-Mpt64 localized to the ER in both yeast and mammalian cells. Additionally, endogenous Mpt64 localized to the ER during Mtb infection of macrophages, suggesting that the observed localization of Mpt64 is not an artifact of heterologous over-expression. Mpt64 did not co-localize with the ER after infection with a Type VII secretion system mutant underscoring the importance of the membrane disrupting properties of ESAT-6 in establishing communication with the host cell (99, 100). This ESX-1 dependent mechanism of cytoplasmic access is similar to the route taken by the autotransporter-like protein tuberculosis necrotizing toxin (TNT) (101, 102). Thus, our data strengthen the argument that the Type VII secretion system facilitates access of non-ESX-1 substrates beyond the phagosome and into the host cell.

We identified PI4P and PI5P as primary monophosphotidylinositol targets of recombinant Mpt64, in addition to the di- and tri-phosphotidylinositols PI(4,5)P_2_ and PI(3,4,5)P_3_. While PI4P is thought to be enriched in the Golgi (103), it also has an established role in mediating protein trafficking from ER exit sites (104, 105). Thus, the ability of Mpt64 to block secretion may stem from its subcellular localization at the ER, interaction with PI4P, and interference with ER to Golgi trafficking. Less is known about PI5P as its basal level is only about 1% of PI4P (106). However, PI5P is increased during bacterial infection and other stresses, and can be found throughout the cell, including the ER (106). Likewise, while PI(4,5)P_2_, the most abundant phosphotidylinositol, is distributed predominantly at the plasma membrane, it can also be found within the cell at the Golgi and ER (107), and PI(4,5)P_2_ serves as the precursor for cellular production of PI(3,4,5)P_3_ (108). Taken together, while Mpt64 binds multiple phosphotidylinositols in vitro, the relative contributions of binding to individual molecules and the role of such binding in mediating the host secretion blockade of Mpt64 remain unknown.

Disruption of the RD2 locus in Mtb H37Rv leads to decreased bacterial burdens in the lungs and spleen of aerosol-infected mice at 3 weeks after infection (80). As Mpt64 is within the RD2 locus, we hypothesized that the single, in-frame deletion of Mpt64 might explain the attenuation phenotype of the RD2 mutant. In line with this prediction, we observed decreased bacterial burdens of MtbΔMpt64 compared to WT Mtb in the lungs of mice at 3 weeks post-infection. Although this decrease was not statistically significant and was not associated with a survival defect, it does suggest Mpt64 contributes to the virulence of the RD2 region. Other genes located in the RD2 locus that were not complemented in the RD2 survival study (80) such PE_PGRS35 (Rv1983) and cfp21 (Rv1984) may also contribute alongside Mpt64 to the virulence defect observed in RD2 deletions. Furthermore, it is possible that one or more of the other Mtb secreted proteins we identified, including the 27 proteins that also localize to the ER are able to perform a redundant function to that of Mpt64 during an animal infection. In a similar vein, whereas *L. pneumophila* encodes over 300 effectors, individual *L. pneumophila* effector deletion mutants are not defective for growth in cells or mice (109, 110). Thus, future work disrupting multiple Mtb MEPs simultaneously will help address the issue of redundancy.

When we infected mice with MtbΔMpt64::Mpt64-NS, a strain of Mtb that still expresses Mpt64 but cannot secrete it into the host cell, we recovered significantly fewer CFU compared to WT from the lungs of MtbΔMpt64::Mpt64-NS infected mice. We hypothesize that this strain suffers from two detrimental consequences. First, blocking Mpt64 secretion prevents it from exerting its virulence function in the host. Second, non-secreted Mpt64 can still be cross-presented to the adaptive immune system (111), thus leading to a cell mediated immune response against Mpt64. This observation is consistent with data that both human patients with active tuberculosis and their PPD positive contacts have T-cell responses to Mpt64 (112) and T-cell reactive Mpt64 epitopes have been mapped (113). Furthermore, Mpt64 staining is observed in granulomas of infected individuals (114, 115). Thus, Mpt64 is highly immunogenic during human infection with Mtb and suggests an evolutionary tradeoff between the virulence function of Mpt64 and its antigenicity. In fact, when we explored the importance of Mpt64 in human disease, we observed that Mpt64 secreted from wild type bacteria localized to the ER of infected human monocyte-derived macrophages and its presence enhanced the survival of Mtb within primary human macrophages. Together these findings suggest that Mpt64 contributes to the virulence of Mtb in human disease.

## Materials and Methods

### Bacterial Strains and Growth Conditions

*M. tuberculosis* Erdman and mutants were grown in Middlebrook 7H9 broth or on Middlebrook 7H11 agar (Difco) supplemented with 10% oleic acid-albumin-dextrose-catalase (OADC, Remel). Liquid medium was also supplemented with 0.05% Tween 80.

### Yeast Strains and Assays

The *Saccharomyces cerevisiae* strain INVSc1 (Invitrogen) was grown at 30°C in histidine drop out media (SD/-HIS) or agar plates (Clontech). The construction of cdc25^ts^ was previously described (68). cdc25^ts^ was grown at 25°C in leucine drop out media or agar plates (SD/-LEU) (Clontech).

The cdc25^ts^ strain and INVSc1 strain were transformed using a lithium acetate (LiAc) protocol. Yeast were grown to high density overnight at the appropriate temperature, shaking. The cultures were diluted to an OD_600_=0.2 in 50mL YPD and allowed to reach mid-log phase. Cells were washed, resuspended in 0.1M LiAc and incubated 10 minutes at room temperature. The sample DNA was mixed with an equal volume of pre-boiled Yeastmaker Carrier DNA (Clontech). To the DNA was added 100uL yeast and 500uL of a solution of LiAc+PEG (40% PEG w/v, 0.1M LiAc). This solution was incubated 30 minutes at 25°C (cdc25^ts^) or 30°C (INVSc1) with agitation every 10 minutes. DMSO was added and the cells were heat shocked at 42°C for 15 minutes. The cells were pelleted, washed in TE and resuspended in 500uL TE. The transformed cells were plated on selective agar plates and incubated at the appropriate temperature for 2-4 days.

To perform the Ras rescue assay, 3-4 fresh colonies were combined in 30μL SD/-LEU and 3 μl was spotted onto duplicate plates that were subsequently incubated at either 25°C or at 37°C for 2 days.

INVSc1 yeast were transformed with a galactose-inducible vector (p413GALGFP) containing GFP-Mtb fusion proteins and selected on SD/-HIS. To induce protein expression, yeast were inoculated in 3mL galactose/raffinose (Gal/Raf) base lacking histidine (Clontech) and allowed to grow 16-20 hours at 30°C, shaking. Cultures were pelleted, resuspended in 30-50uL PBS and immobilized on an agar pad prior to visualization.

### Yeast Lysis and Western Blotting

Yeast (cdc25^ts^) were inoculated into 5mL SD/-Leu and incubated overnight at room temperature, shaking (250rpm). To lyse, 1.5mL of each culture was centrifuged at 14,000rpm for one minute. Each pellet was resuspended in 100μL 2.0M LiAc and incubated on ice for five minutes. Samples were centrifuged at 14,000 for one minute to pellet, resuspended in 100μL 0.4M NaOH and incubated on ice for five minutes. Samples were pelleted as before, resuspended in 75μL 1 × SDS Laemmli sample buffer and boiled at 100°C for five minutes. Lysates were centrifuged 14,000rpm for one minute to remove to debris, separated by SDS-polyacrylamide gel electrophoresis and transferred to polyvinylidene difluoride membrane for Western blotting. Fusion proteins were detected by rabbit anti-Ras (1:100) and equal loading was confirmed by detection with rabbit anti-G6PDH (1:10,000).

### Cell Culture

HeLa cells were cultured in Dulbecco’s modified Eagle medium (DMEM, Gibco) supplemented with 10% fetal bovine serum (FBS, Gibco), 100 I.U./mL penicillin, 100 μg/mL streptomycin, 292 μg/mL L-glutamine (Corning). RAW267.4 macrophages were cultured in RPMI 1640 (Gibco) supplemented with 10% heat-inactivated FBS, 100 I.U./mL penicillin, 100 μg/mL streptomycin, 292 μg/mL L-glutamine, and 10mM HEPES (Hyclone).

Primary human macrophages were isolated from buffy coats from anonymous donors provided by a local blood bank. To isolate, 50mL of blood from each donor was added to an equal volume of PBS then separated by centrifugation over a Ficoll-Paque Plus gradient at 750 × g for 20 minutes with no brake. The lymphocyte/monocyte layer was collected and incubated 1-2 minutes with 1mL ACK lysing buffer (Gibco) to remove red blood cells. The cells were diluted to 50mL in PBS and centrifuged 350 × g for 10 minutes at 4°C. The supernatant was removed and cells were washed in 25mL PBS and pelleted at 160 x g for 15 minutes at 4°C. Cells were washed again in 25 mL PBS but centrifuged at 300 × g for 10 minutes at 4°C. This final pellet was resuspended in 5-10 mL of RPMI 1640 supplemented with 10% human AB serum (Corning). To differentiate into macrophages, cells were cultured in RPMI 1640 supplemented with 10% human AB serum and 50ng/mL human M-CSF (R&D Systems) for at least 4 hours, washed in PBS then replaced with RPMI 1640 +10% human AB serum + human M-CSF (50ng/mL) for 7 days with media changes every 1-2 days.

### Antibodies

To generate an antibody against native Mpt64, two rabbits were immunized with recombinant 6 × HIS-tagged Mpt64ΔSP purified from *E. coli* in incomplete Freund’s adjuvant (Pacific Biosciences). The polyclonal rabbit antibody to Antigen 85 and mouse anti-GroEL2 (CS-44) are from BEI Resources. Chicken, mouse and rabbit anti-calreticulin were purchased from Abcam and anti-GM130 was purchased from BD Biosciences. Mouse anti-Tom20 F-10 was purchased from Santa Cruz. Rabbit anti-Ras and rabbit anti-glucose-6-phosphate dehydrogenase (G6PDH) were purchased from Cell Signaling Technology and Sigma, respectively. Mouse anti-PMP70 and horse radish peroxidase (HRP)-conjugated secondary antibodies were purchased from Thermo Scientific. Alexa fluor-conjugated secondary antibodies were from Life Technologies.

### Molecular Biology

Unless otherwise stated, all Mtb proteins were cloned from the BEI resources Gateway Mtb ORF library using Gateway cloning technology (Life Technologies). The Mpt64 truncation mutants were PCR amplified (Supplemental Table 2) and cloned into pENTR (Life Technologies) prior to cloning into subsequent destination vectors.

### hGH Secretion Assay and Quantification

HeLa cells were plated in 24-well plates to achieve approximately 50,000 cells/well 24h prior to transfection. Cells were co-transfected with 1 μg hGH-CAD and 1μg GFP-Mtb effector or GFP alone using FuGene 6 (Promega) per manufacturer instructions. Cells were transfected 16-18h at 37°C 5% CO_2_. The transfection media was then aspirated and replaced with DMEM containing 2 μM D/D Solubilizer (Clontech) and incubated for 2 h at 37°C 5% CO_2_. The plates were centrifuged at 1500 RPM for 5 minutes to pellet debris and the culture supernatants were saved at −80°C prior to hGH quantification.

Secreted hGH was quantified by ELISA (Roche, 11585878001). Briefly, samples were thawed on ice and 20 μL was transferred to each well containing 180μL sample buffer (1:10). The plate was incubated 1h at 37°C, washed 5 times in 250μL wash buffer and incubated 1h at 37°C with a polyclonal antibody to hGH conjugated to digoxigenin (α-hGH-DIG). The plate was washed as described and incubated 1 h at 37°C with a polyclonal antibody to digoxigenin conjugated to peroxidase (α-DIG-POD). The plate was washed and developed in peroxidase substrate (2,2′-Azinobis [3-ethylbenzothiazoline-6-sulfonic acid]-diammonium salt). The absorbance was read on a Biotek plate reader at 405nm.

### PIP strips membrane binding

6xHIS-Mpt64_24-228 and 6xHIS-Mpt64_24-143 were purified by cobalt TALON affinity resin (Clontech). PIP strips (Invitrogen) were blocked for one hour at room temperature in 3% fatty-acid free bovine serum albumin (BSA) in TBST. Mpt64_24-228 or Mpt64_24-143 was diluted to 1.5ug/mL in 3mL 3% fatty-acid free BSA and incubated with the PIP strips for 3h at room temperature with agitation. Membranes were washed three times in 3% fatty-acid free BSA prior to incubation with anti-Mpt64 or pre-immune serum (1:3,000) overnight at 4°C, with agitation. Membranes were washed three times in 3% fatty-acid free BSA then incubated with HRP-conjugated donkey anti-rabbit (1:2000) for 30 minutes at room temperature. Membranes were washed three times before detection of Mpt64 lipid interactions by chemiluminescence.

### Transfection and co-localzation of MEPs in HeLa cells

HeLa cells were transfected overnight with GFP fusion proteins using FuGene 6 transfection reagent (Roche). Cells were fixed in 4% paraformaldehyde (PFA) for 15 minutes, washed in PBS and permeabilized in 0.25% Triton X-100 for 3 minutes at room temperature or in 100% methanol for 10 minutes at −20°C when using Tom20 antibody. Cells were stained with organelle-specific antibodies for 1h at room temperature. Antibodies were visualized by secondary antibodies conjugated to Alexa Fluor 594. Cells were mounted in ProLong Gold + DAPI and z-stacks were collected on an AxioImager M2 microscope (Zeiss).

### Infection and co-localization of Mpt64 in macrophages

Bacteria were washed repeatedly in PBS, then sonicated to create a single cell suspension. RAW267.4 cells were infected in DMEM+10% horse serum at MOI 20:1 with mycobacteria expressing mCherry. Cells were centrifuged at 1500 rpm for 10 minutes to permit bacterial attachment, then allowed to phagocytose for 1.5h at 37°C 5% CO_2_. Cells were fixed after 4 and 24 hours post-infection in 4% PFA for 60 mintues Cells were permeablized in 0.25% Triton X-100 for 3 minutes at room temperature then blocked in 5% normal donkey serum (Sigma). Mpt64 was detected with rabbit anti-Mpt64 antibody (1:500) and an HRP-conjugated goat-anti rabbit secondary antibody (1:1000, Santa Cruz). Antibody signal was amplified by addition of biotinylated Tyramide (1:50, PerkinElmer) with detection by Alexa fluor 488-conjugated streptavidin (1:250, Jackson Immunoresearch) or Cyanine 5 Tyramide (1:50, PerkinElmer). Z-stack slices were acquired with an AxioImager M2 microscope (Zeiss).

Primary human macrophages were infected in RPMI + 10% human AB serum at an MOI 10:1 with mycobacteria expressing mCherry for 2h at 37°C 5% CO_2_ to allow for phagocytosis. Cells were washed and fixed at 4 hours post-infection in 4% PFA for 45-60 minutes. Cells were permeabilized in 100% ice cold methanol for 10 minutes at - 20°C and blocked in 5% normal goat serum (Sigma). Mpt64 was detected with rabbit anti-Mpt64 antibody (1:500) and an HRP-conjugated donkey-anti rabbit secondary antibody (1:500, Thermo Scientific) followed by amplification with cyanine 5 Tyramide (1:50, PerkinElmer). Co-localization of Mpt64 with the ER was detected with chicken anti-calreticulin (1:100), followed by goat anti-chicken-488 (Abcam).

### Macrophage infections for CFU

Primary human macrophages were seeded in low-evaporation 24-well plates at approximately 5 × 10^5^ cells/well. Bacteria were washed repeatedly in PBS, then sonicated to create a single cell suspension. Macrophages were infected in RPMI + 10% human AB serum at MOI 0.5 for 2 hours at 37°C 5% CO_2_ to allow phagocytosis to occur. The cells were washed in PBS then replaced with RPMI + 10% human AB serum and cells were washed every day between time points. The cells were lysed at time zero and subsequent time points in 500 μL 0.5% Triton X-100 in PBS. Serial dilutions were plated on 7H11 plates and colonies were enumerated after 2-3 weeks.

### Construction of the Mtb Mpt64 deletion mutant and complementation

An in-frame Mpt64 deletion in Mtb was made using mycobacteriophage as previously described (92). Briefly, 500-bp 5’ to the Mpt64 start codon and 500-bp 3’ to the Mpt64 stop codon were amplified from Erdman genomic DNA (Supplemental Table 2) and sequentially cloned into the multiple cloning sites of pMSG360HYG. This vector was linearized with AflII and DraI and transformed into EL350/phAE87 *E. coli* by electroporation. Phagemid DNA was isolated from pooled colonies and transformed into *M. smegmatis* by electroporation. Plaques were isolated and pooled from *M. smegmatis* lawns and high titer phage was produced. Log phase Mtb Erdman was transduced with phage at 42°C for 4h. Mutants were selected on 7H11+Hygromycin (100μg/mL). Wild-type Mpt64 and Mpt64 lacking its secretion signal were cloned into an integrating vector containing a constitutive promoter (pMV306_MSP), conferring zeocin resistance. The MtbΔMpt64 was transformed by electroporation and complements were selected on 7H11 + zeocin (25 μg /mL).

To confirm expression and secretion of Mpt64 complements, Mtb strains were grown to late-log phase and pelleted by centrifugation. The culture supernatants were saved and passed twice through 0.22 μm filters. Bacterial pellets were boiled 30 minutes in lysis buffer (50mM Tris, pH 7.4, 150mM NaCl) supplemented with Complete Mini protease inhibitor (Roche), then subjected to bead beating to lyse the cells. Protein content in lysates was determined by Bradford assay. Mpt64 expression in the lysates and culture supernatants was detected by Western blotting using a rabbit polyclonal antibody to Mpt64 (1:10,000). Equal loading of samples in the lysates and supernatants was confirmed by Western blotting with anti-GroEL2 (1:500) and anti-antigen 85 (1:1000) respectively.

### Mouse infections

Female BALBc mice (The Jackson Laboratory) were infected via aerosol as described previously (116). Briefly, mid-log phase Mtb were washed in PBS repeatedly then sonicated to create a single-cell suspension. Bacteria were resuspended to yield an OD_600_=0.1 in PBS. This suspension was transferred to the nebulizer of a GlassCol aerosolization chamber calibrated to infect mice with ∼100 bacteria per animal. On the day of infection, whole lungs were collected from 5 mice per group, homogenized and plated on 7H11 to determine initial inoculum. At subsequent time points, the left lung, spleen and left lobe of the liver were used to determine CFU, while the right lung was insuflated with 10% neutral buffered formalin for histopathology. Animal experiments were reviewed and approved by the Institutional Animal Care and Use Committee at the University of Texas Southwestern.

### Statistical analysis

Statistical analysis was performed using GraphPad Prism software. For in vivo CFU calculations and area of lung inflammation, the non-parametric Kruskal-Wallis test with Dunn’s multiple comparison was used. Analysis of survival studies was performed by Kaplan-Meier test. In vitro fold CFU was calculated by dividing the average CFU of quadruplicates of each Mtb strain at each time point by the average at Day 0. Propagation of error (z) for each fold calculation was determined by the equation:

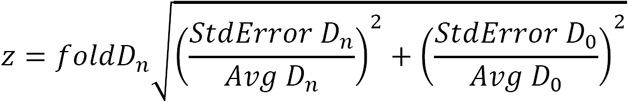

where D_n_ represents the time point value for any given strain and D_0_ represents the day zero values for that given strain (117). Statistical analysis on fold CFU was performed by ANOVA.

### Lysozyme pull down

E. coli lysates containing 6x-histidine (6xHIS) tagged Mpt64 or an unrelated protein Cor were incubated with cobalt affinity resin (TALON, Clontech) to bind histidine-tagged proteins. After extensive washing, 1mg/mL either hen egg white or human lysozyme was flowed over the immobilized beads and incubated 5 minutes. Beads were washed two more times before proteins were eluted with 300mM imidazole.

### Mtb genomic DNA isolation

Late exponential phase Mtb was collected by centrifugation and washed once in PBS. Pellets were boiled 20-30 minutes to sterilize. Pellets were washed once in GTE (25mM Tris, pH 8.0; 10mM EDTA; 50mM glucose) and incubated overnight in lysozyme solution (10mg/mL in GTE) at 37°C. Samples were incubated in 10% SDS and 10mg/mL proteinase K for 40minutes at 55°C followed by incubation in NaCl and CTAB (2.4M NaCl, 274mM cetrimonium bromide (Sigma)) at 60°C for 10 minutes. Genomic DNA was then isolated using a phenol-chloroform extraction followed by ethanol precipitation.

### Extraction of apolar lipids and PDIM analysis

Log phage Mtb or MtbΔMpt64 were synchronized to OD_600_=0.2 in 7H9 supplemented with 0.01% Tween 80 and grown 24 hours. Bacteria were collected by centrifugation at 1,600xg for 10 minutes, resuspended in 1mL 15% isopropanol and transferred to a glass tube containing 5mL chloroform: methanol (17:1, v/v) and incubated 24 hours at room temperature. Samples were centrifuged at 1,600xg for five minutes and the apolar lipids were collected from the bottom, organic layer and dried. Apolar lipids were resuspended in 1.5mL 100% methanol. Tween 80 was removed by addition of cobalt/thiocyanate solution and vortexed. Remaining lipids were extracted by addition of 4mL hexane. After centrifugation the organic layer was saved and the aqueous layer was re-extracted with 4mL hexane. Both hexane fractions were combined, dried and resuspended in 1mL chloroform: methanol (2:1, v/v). PDIM standard was similarly resuspended. PDIM standard, or apolar lipids extracted from Mtb or MtbΔMpt64 were infused into an AbSciex TripleTOF 5600/5600+ mass spectrometer. Samples were analyzed in the positive mode.

## Acknowledgements

We would like to thank Patrick Cherry and Molly Moehlman for their contributions to this work during their summer internships. We thank BEI resources for providing reagents. This work was supported by NIH Grants R01 AI099439, R21 AI111023 and U01 AI125939 (to M.U.S.) and T32AI007520 (to C.E.S and B.L.P). M.U.S. is a Disease-Oriented Clinical Scholar at University of Texas Southwestern Medical Center.

C.E.S., N.M.A. and M.U.S. conceived and designed experiments. B.A.W. provided reagents and experimental advice. C.E.S., and S.C. performed immunofluorescence microscopy. C.E.S., B.L.P., L.H.F., and V.R.N. performed mouse aerosol infections with Mtb. B.L.P. performed mass spectrometry analysis on PDIM. C.E.S. performed the remaining research include screening experiments, molecular biology, bacterial genetics and macrophage infections. C.E.S. and M.U.S. drafted the manuscript. All authors edited and approved of the final manuscript.

---

Supplemental Figure 1. Recombinant Mpt64 does not interact with lysozyme. (**A**) Sequence and secondary structure alignment of Mpt64 and *B. subtilis* RsiV (template) performed by Phyre^2^ software (http://www.sbg.bio.ic.ac.uk/phyre2/). The Mpt64 DUF3298 is indicated by a red line above the Mpt64 amino acid numbering. (**B**) Mpt64 (M) or an unrelated protein Cor (C) were immobilized on cobalt beads and hen egg white (HEW) or human lysozyme (HuLYZ) was incubated with these or beads alone (B) for five minutes. After washes, proteins were eluted with 300mM imidazole.

Supplemental Figure 2. Construction and phenotypic analysis of MtbΔMpt64. (**A**) Western blot detecting expression of Mpt64, Ag85 and GroEL2 in either the lysate of the cell pellet (P) or culture supernatant (S) of wild type Mtb or MtbΔeccD1. (**B**) Detection of hygromycin resistance cassette insertion in place of *mpt64*. Genomic DNA from wild type Mtb or MtbΔMpt64 was amplified by polymerase chain reaction and products were analyzed by agarose gel electrophoresis. (**C**) Growth of Mtb, MtbΔMpt64, MtbΔMpt64::Mpt64 and MtbΔMpt64::Mpt64-NS in 7H9 measured by optical density at 600nm. (**D**) PDIM standard or apolar lipid extracts from Mtb or MtbΔMpt64 were analyzed on an AbSciex TripleTOF 5600/5600+ mass spectrometer.

Supplemental Figure 3. Secreted Mpt64 co-localizes with calreticulin in murine macrophages. (A) RAW267.4 murine macrophages were infected with the indicated strains of mCherry expressing Mtb (cyan) for four hours and subsequently stained for Mpt64 (red) and calreticulin (green). Nuclei are stained in blue. Scale bars are 10μm.

Supplemental Figure 4. Secreted Mpt64 co-localizes with calreticulin in human macrophages. (A) Primary human macrophages were infected with the indicated strains of mCherry expressing Mtb (cyan) or left uninfected for four hours prior to fixation and staining for Mpt64 (red) and calreticulin (green). Scale bars are 5μm.

